# Response of *Escherichia coli* chemotaxis pathway to pyrimidine deoxyribonucleosides

**DOI:** 10.1101/2025.04.13.648572

**Authors:** Malay Shah, Victor Sourjik

## Abstract

Nucleosides are essential components of all living cells. Bacteria use salvage pathways to import nucleosides from their environment and to utilize them for nucleic acid biosynthesis, but also as alternative sources of carbon, nitrogen and energy. Motile bacteria commonly show chemoattraction towards nutritionally valuable compounds, and in this work, we demonstrate that the chemotaxis pathway of *Escherichia coli* exhibits specific attractant response to pyrimidine nucleosides. Particularly sensitive response, in the sub-micromolar range, was observed for deoxyribonucleosides thymidine and deoxycytidine. In contrast, ribonucleosides elicited weaker and less sensitive response, and no response to pyrimidine nucleobases was observed in the micromolar range of concentrations. Our subsequent analysis revealed that the pathway response to pyrimidine deoxyribonucleosides is mediated by the minor *E. coli* chemoreceptor Tap. The observed narrow dynamic range of this response indicates that sensing of deoxyribonucleosides is indirect, likely via an unknown periplasmic binding protein that interacts with Tap.

## Introduction

Motile bacteria use chemotaxis to navigate chemical gradients in their environment, locating optimal niches for their proliferation and survival (1-3). The gradient navigation relies on the detection of temporal changes in ligand concentrations, to which bacteria respond by adjusting their rate of tumbling. The molecular details of this process are well understood (1, 4). Ligands are typically bound by the transmembrane chemoreceptors that form a ternary complex with an adaptor protein CheW and a histidine kinase CheA. Attractant binding induces a conformational change that inactivates the autophosphorylation activity of CheA, thereby lowering phosphorylation of the response regulator CheY. Since phosphorylated CheY (CheY-P) interacts with the flagellar motor to induce cell tumbles, reduced levels of CheY phosphorylation result in prolonged periods of straight swimming up the gradients of attractants. In addition to this, chemotaxis machinery of *E. coli* includes the CheY-P phosphatase CheZ, as well as the adaptation module consisting of CheR and CheB that modulates receptor activity and sensitivity to ligands through methylation and demethylation on specific glutamate residues.

The chemotaxis pathway in *E. coli* is known to sense a number of nutrient compounds, including amino acids (5, 6), sugars (7, 8) and dipeptides (9), but also signalling molecules such as indole (10-12), autoinducer 2 (11, 13, 14) and mammalian hormones (10, 15, 16). *E. coli* has four transmembrane chemoreceptors: two major receptors, Tar and Tsr, and two minor receptors, Trg and Tap (1, 4). Primary ligands of Tar and Tsr are amino acids aspartate and serine, respectively, that bind directly to their periplasmic sensory domains. Other chemoeffectors are sensed through the interaction between the receptor sensory domains and a ligand-bound periplasmic binding protein that is a component of an ATP binding cassette (ABC) transporter for that ligand. Such indirect sensing has been established for maltose in the case of Tar (7), and for the autoinducer 2 in the case of Tsr (13). Indirect sensing is further the only established more of responses mediated by minor receptors, including responses of Trg to glucose, galactose and ribose (17), and of Tap to dipeptides (9). Besides these well-established mechanisms, cytoplasmic regions of chemoreceptors can mediate responses to unconventional stimuli such as phenol, temperature, or cytoplasmic pH (18), as well as to the substrates of the phosphotransferase system (PTS) (8), although these sensory modes remain poorly understood.

Amino acids and sugars are known chemoeffectors not only for *E. coli* but also for other bacteria, supporting the notion that bacteria are attracted by nutrients (19). Another common class of bacterial receptor ligands are nucleobases and their derivatives (20-23). Indeed, like other bacteria, *E. coli* can uptake and salvage nucleobases and nucleosides from the medium to replenish its nucleotide pool but also to use them as alternative sources of carbon, nitrogen and energy (24). A previous study has shown that *E. coli* exhibits a Tap-mediated chemoattraction to millimolar concentrations of pyrimidine nucleobases (25), but the mechanism and scope of this response was not further investigated. Here, we demonstrate that Tap mediates highly sensitive response to pyrimidine deoxyribonucleosides, and lower-affinity response to pyrimidine ribonucleosides, whereas no response to pyrimidine nucleobases thymine or cytosine was observed in the tested micromolar concentration range. Our results further suggest that this sensing of nucleosides by Tap is indirect, likely mediated by a yet unknown periplasmic binding protein.

## Results

### Characterization of *E. coli* chemotaxis pathway response to nucleosides

To systematically characterize the response of *E. coli* chemotaxis pathway to pyrimidine nucleobases, ribonucleosides and deoxyribonucleosides, we used an assay of the pathway activity based on the Förster resonance energy transfer (FRET) (26). For that, we expressed the pair of interacting pathway proteins fused to cyan and yellow fluorescent proteins, CheZ-CFP and CheY-YFP. Since the interaction of CheY with its phosphatase depends on phosphorylation, it allows real-time monitoring of the pathway activity in a cell population using FRET (26, 27). To test their response to different pyrimidines, *E. coli* cells expressing the FRET pair were immobilized in the flow chamber and exposed to different concentrations of tested compounds. FRET responses were recorded as a change in ratio of the YFP to CFP fluorescence (FRET ratio), with L-aspartate, a canonical chemoattractant for *E. coli*, being used as a positive control. Since attractant inhibits CheA activity, leading to a reduction in the fraction of interacting CheY-YFP and CheZ-CFP proteins, addition of L-aspartate caused a rapid decrease in the FRET ratio (Fig. 1A). The response to 100 μM aspartate is known to completely inhibit the pathway activity, thus reducing the YFP to CFP ratio to the level observed in the absence of energy transfer (27). Subsequent removal of an attractant leads to an increase in the CheA activity with concomitant increase in the fraction of interacting FRET reporter proteins. The resulting pathway activity is initially higher than that in the adapted cells, because the CheR-mediated adaptive changes in receptor methylation during prolonged stimulation with attractant increase receptor activity bias. This transient overshoot of the pathway activity upon removal of the chemoattractant is gradually reset to the steady-state level by the CheB-dependent receptor demethylation.

**FIG 1.**
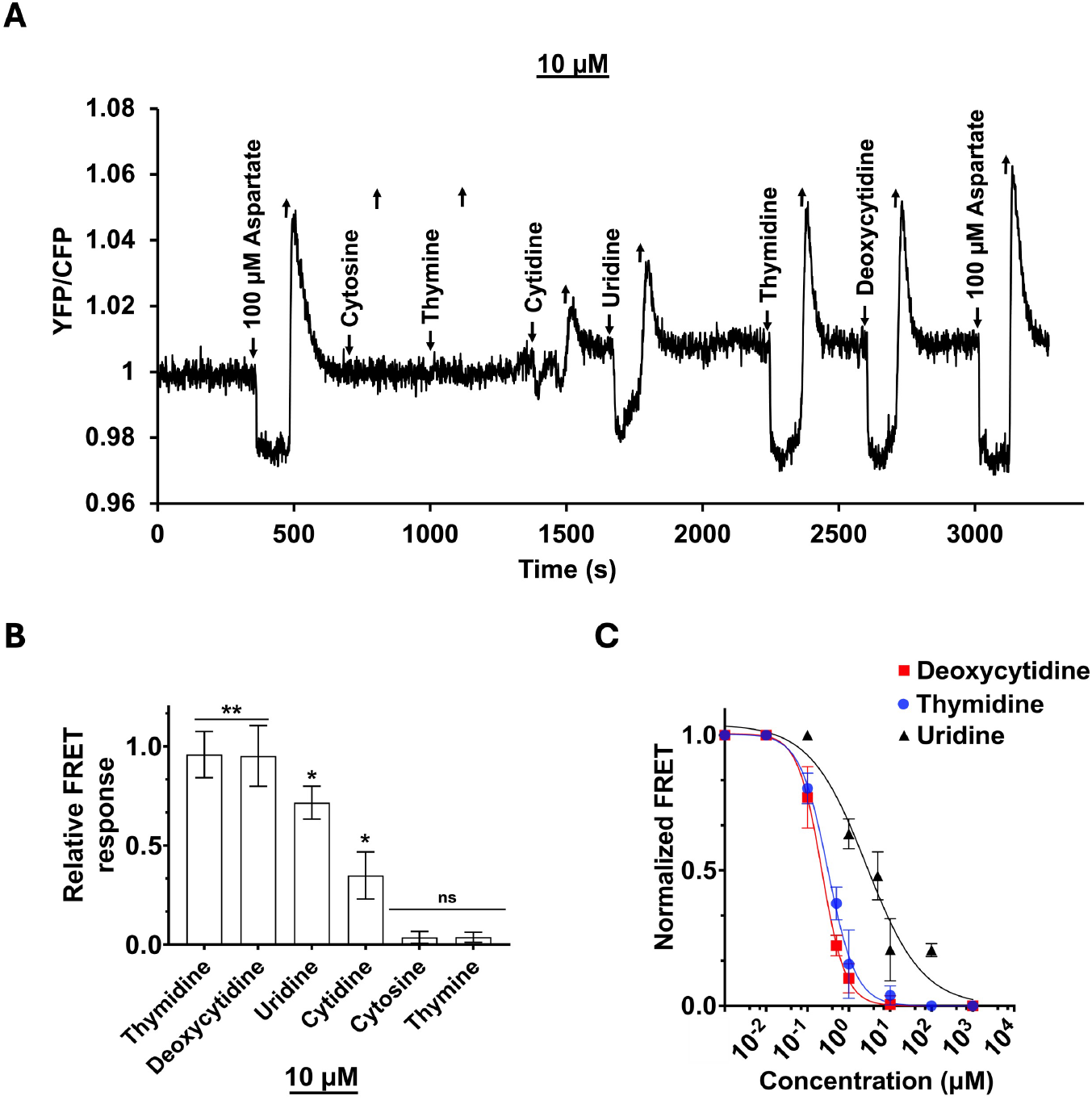
FRET measurement of the chemotaxis pathway response to pyrimidine nucleosides and nucleobases. (A) *E. coli* cells expressing CheZ-CFP/CheY-YFP FRET pair were immobilized in a flow chamber, adapted to tethering buffer and stimulated by addition and subsequent removal of 10 μM of indicated compounds. Saturating stimulation with 100 μM L-aspartate was used as a control. FRET measurements for all figures were detrended as described in the methods. FRET signal is represented by the ratio of YFP to CFP emission, normalized to the YFP/CFP ratio at the beginning of the measurement and compensated for the gradual drift in the baseline as described in the Material and methods. (B) Response amplitudes of buffer-adapted cells to 10 μM of indicated compounds, normalized in each experiment to the response amplitude for 100 μM L-aspartate, and calculated from 3 biological replicates. Error bars indicate standard deviations. Significance of the response compared to zero was assessed using Student’s one-sample *t*-test (ns: non-significant, \*\**P*-value≤0.01, \**P*-value≤0.05). (C) Dose-response curves measured by FRET for indicated compounds. Cells were stimulated by step-like addition and subsequent removal of increasing amounts of an attractant. After each stimulation cells were re-adapted to the buffer. The FRET response for each step was normalized to the maximal FRET response. Error bars indicate standard deviations and means are derived from 3 biological replicates. Data were fit using the Hill equation, with EC_50_ fit values and respective 95 % CI (shown in brackets) for each compound being 3.7 μM (2.3 μM to 5.6 μM) for as uridine; 237 nM (200 nM to 280 nM) for deoxycytidine; 337 nM (277 nM to 400 nM) for thymidine.

Our results demonstrate that, in the tested concentration range of 1 to 100 μM, stimulation with pyrimidine nucleobases (cytosine and thymine) displayed no measurable FRET response (Fig. 1A,B and Fig. S1A,C). In contrast, ribonucleosides (cytidine and uridine) elicited a distinct attractant response, although it was only observed at high concentrations (10 and 100 μM) and weaker than the response to aspartate (Fig. 1A,B and Fig. S1B,D). The strongest and most sensitive response was observed upon stimulation with deoxyribonucleosides (deoxycytidine and thymidine), with a distinct change in FRET already at 1 μM (Fig. S1B) and the amplitude of response being similar to stimulation with aspartate at 10 μM.

To further compare the sensitivity of *E. coli* chemotaxis pathway to pyrimidine deoxyribonucleosides and ribonucleosides, we measured the dose-dependence of the FRET response to deoxycytidine, thymidine and uridine (Fig. 1C). Whereas the half-maximal inhibition of the pathway activity by uridine was observed at EC_50_ of ∼4 μM, the EC_50_ values for deoxycytidine and thymidine were tenfold lower, at ∼300 nM. Such sensitivity is within the same range as for the established high-affinity *E. coli* attractants, including aspartate and serine that bind to their respective chemoreceptors directly, and galactose, ribose or dipeptides that are sensed via periplasmic binding proteins (28). We thus conclude that pyrimidine deoxyribonucleosides are high-affinity chemoattractants for *E. coli*.

### Sensing of pyrimidine nucleosides is likely indirect and mediated by Tap

To investigate whether the sensing of pyrimidine deoxyribonucleosides is direct or indirect, we next investigated the dynamic range of the chemotaxis pathway response. The dynamic range represents the most characteristic difference between chemoeffector ligands that directly bind to *E. coli* chemoreceptors, and those that rely on binding to periplasmic proteins to relay the signal to chemoreceptors (28). Directly binding ligands can be sensed over four to five orders of magnitude of ambient concentrations (5, 27), because receptor methylation system enables efficient tuning not only of the pathway activity but also of receptor sensitivity to these ligands as cells become expose to their increasing concentrations. In contrast, while the response to indirectly binding ligands is similarly adaptive in respect to the pathway activity, binding of these ligands to their periplasmic binding proteins saturates and cannot be tuned by the adaptation system. As a consequence, the dynamic range of response to these ligands is much narrower than for directly binding ligands, spanning only two orders of magnitude of ligand concentrations (7, 28).

We thus determined the dynamic range of the pathway response to deoxycytidine and thymidine as done previously (28), by raising concentration of ligands in tenfold steps and allowing cells to adapt prior to each subsequent stimulation. We found that both deoxyribonucleosides show an approximately hundredfold range of concentrations over which they elicit the response upon adaptation, saturating above 10 μM (Fig. 2A and Fig. S2). This range is very similar to those previously observed for indirectly binding ligands galactose, ribose, maltose or dipeptides, and much narrower than the dynamic range for aspartate or serine (28). Thus, we conclude that pyrimidine deoxyribonucleosides are likely sensed indirectly, with the response being mediated by a periplasmic binding protein.

**FIG 2.**
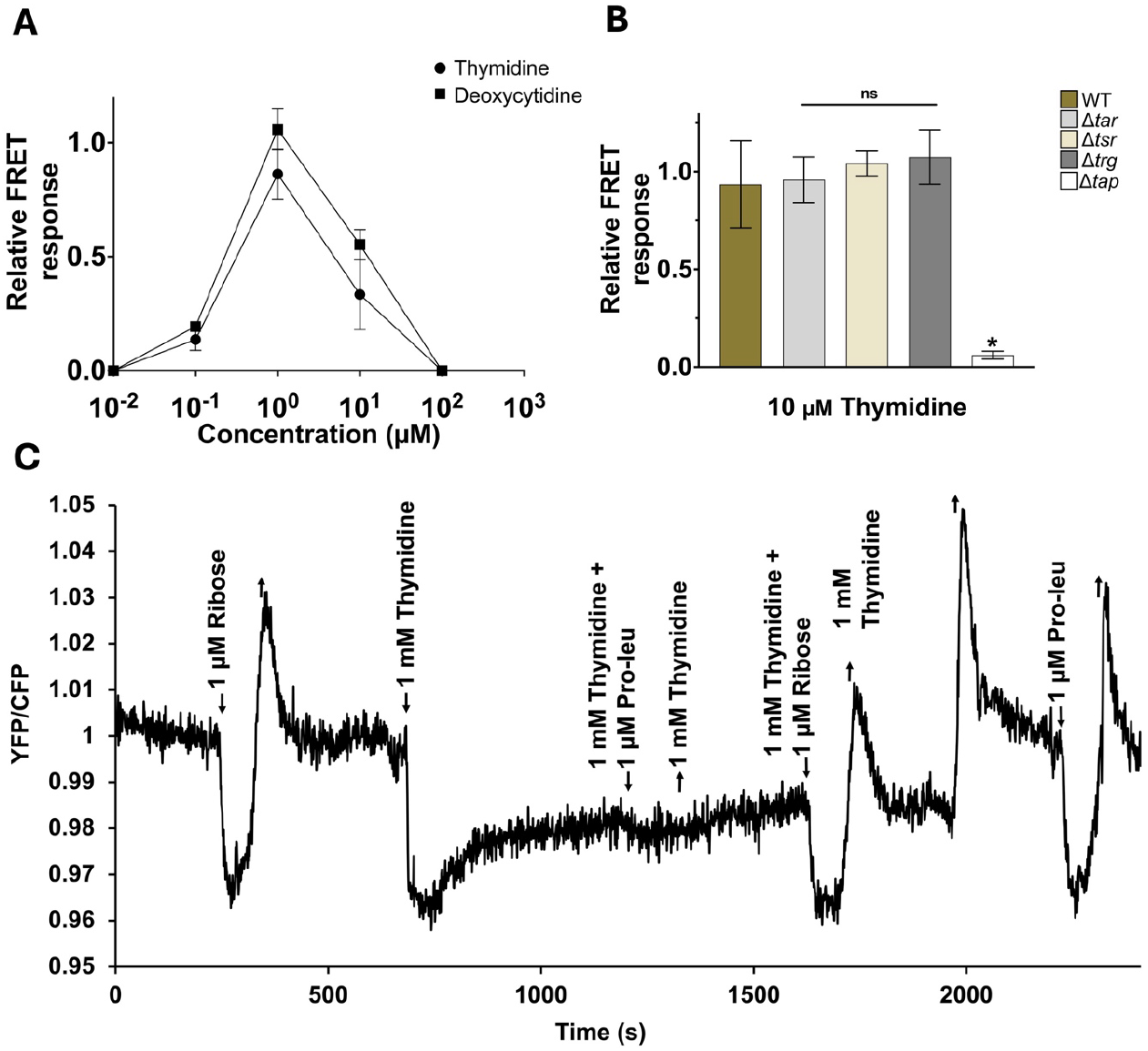
Mechanism of the chemotactic response to pyrimidine deoxyribonucleosides. (A) FRET measurement of the dynamic range of the chemotactic response to deoxyribonucleosides. Concentration was raised in 10-fold steps, and cells were allowed to adapt prior to each subsequent stimulation (see Fig. S2). The response for each step was normalized to the response of buffer-adapted cells towards a saturating stimulus of 100 μM L-aspartate. Experiments were performed in triplicate. Error bars indicate standard deviations. (B) Amplitudes of FRET response to 10 μM thymidine for wild-type *E. coli* and indicated chemoreceptor deletion strains. Each bar represents mean response of 3 biological replicates, normalized to 100 μM L-aspartate or 100 μM Pro-leu (for Δ*tar*). Error bars indicate standard deviations. Statistical significance was calculated using Student’s two-sample *t*-test between wild-type and deletion strain response (ns: non-significant, * *P*-value ≤0.05). (C) Competition between FRET response to thymidine and proline-leucine. Trg ligand ribose was used as control. Buffer adapted cells were stimulated and adapted to 1 mM thymidine and, post adaptation, stimulated with 1 mM thymidine combined with 1 μM proline-leucine (Pro-leu). Stimulation with 1 μM Pro-leu in the absence of thymidine was tested post competition experiment.

To identify the chemoreceptor involved in the sensing of pyrimidine nucleosides, we tested FRET responses of *E. coli* strains deleted for individual chemoreceptor genes. While Δ*tar*, Δ*tsr* and Δ*trg* strains responded to thymidine comparably to the wild type, Δ*tap* strain showed no response to uridine, deoxycytidine or thymidine (Fig. 2B, Fig. S3 and Fig. S4A). This suggests that the observed response to deoxycytidine or thymidine is mediated by Tap. To further confirm it, we performed a competition assay between thymidine and the known ligand of Tap, proline-leucine, in the wild-type cells. Indeed, presence of 1 mM thymidine in the background completely inhibited the FRET response to proline-leucine, while not affecting the response to ribose that is indirectly sensed by another receptor, Trg (Fig. 2C). The response to thymidine was also weakened by the presence of 1 mM proline-leucine in the background (Fig S4B).

## Discussion

Extracellular nucleobases and nucleosides can serve as sources of carbon, nitrogen and energy, or can be used by bacteria to replenish their nucleotide pool (24). Consistent with that, recent studies indicated that chemotaxis to nucleobases and nucleosides may be common among bacteria (19, 21, 22). Surprisingly, however, chemotactic response to these compounds remained little investigated in the model organism *E. coli*. The only previous study on this topic demonstrated that *E. coli* exhibits a Tap-mediated chemoattraction to at millimolar concentrations of pyrimidine nucleobases (25). However, the mechanism and specificity of this response was not further investigated.

Here we could show that Tap mediates much stronger and more sensitive response to pyrimidine deoxyribonucleosides than to pyrimidine ribonucleosides or to nucleobases. Whereas *E. coli* responded to thymidine and deoxycytidine already at sub-micromolar concentrations, the response to cytidine or uridine was observed only about 1 μM, and no response to cytosine or thymine could be observed up to 100 μM. These results suggest that deoxyribonucleosides and not nucleobases are the primary pyrimidine chemoattractants sensed by Tap. The sub-micromolar range of concentrations at which the response to pyrimidine deoxyribonucleosides is observed is similar to the response threshold for the most potent established *E. coli* chemoattractants (28). Pyrimidine deoxyribonucleosides appear to be even stronger effectors for Tap than dipeptides, because the adaptation to thymidine completely abolished the response to the dipeptide proline-leucine, whereas the adaptation to proline-leucine only weakened the response to thymidine.

Similar to dipeptides, pyrimidine deoxyribonucleosides appear to be sensed via the indirect mechanism. This is suggested by the narrow dynamic range of the response to thymidine or deoxycytidine, which is similar to the dynamic range of the response to dipeptides or sugars that are known to be sensed via the periplasmic binding proteins, but very different from the dynamic range of the response to directly binding amino acids (28). Since previous study already showed that the Tap-mediated response to high concentrations of pyrimidine nucleobases does not depend on the dipeptide-binding periplasmic protein DppA (25), it likely relies on another periplasmic protein that remains to be identified.

## Materials and methods

### Bacterial strains and plasmids

*E. coli* W3110 RpoS^+^ (29) was used as the wild-type background for FRET experiments. All chemoreceptor knockout strains were made by P1 transduction from lysates prepared using strains from the Keio collection (30). Km^R^ cassettes were removed by FLP recombination (31). For expression of the FRET pair CheY-YFP and CheZ-CFP, plasmid pVS88 inducible by isopropyl-β-D-thiogalactoside (IPTG) (32) was transformed into respective *E. coli* strains.

### FRET assay

The FRET measurements were performed as previously described (22, 26, 27, 32). Briefly, *E. coli* cells were grown in tryptone broth (TB) media (1% tryptone, 0.5% NaCl), supplemented with 100 mg/mL ampicillin and 50 μM IPTG at 34 °C and 275 r.p.m. Cells were harvested at OD_600_ of 0.6 by centrifugation (4000 × *g* for 5 min), washed with tethering buffer (10 mM KH_2_PO_4_/K_2_HPO_4_, 0.1 mM EDTA, 1 μM methionine, 10 mM sodium lactate, pH 7), and stored at 4 °C for 30 min. For microscopy, the cells were attached to the poly-lysine-coated coverslips for 10 min and mounted into a flow chamber that was maintained under constant flow of 0.3 ml/min of tethering buffer using a syringe pump (Harvard Apparatus) that was also used to add or remove compounds of interest. FRET measurements were performed on an upright fluorescence microscope (Zeiss AxioImager.Z1) equipped with photon counters (Hamamatsu). CFP fluorescence was excited at 436/20 nm through a 455 nm dichroic mirror by a 75 W Xenon lamp. To detect CFP and YFP emissions, 480/40 nm band pass and 520 nm long pass emission filters were used, respectively. Fluorescence of a monolayer of 300–500 cells was continuously recorded in the cyan and yellow channels using photon counters with a 1.0 s integration time. The fluorescence signals were analysed as described previously (26).

Gradual drift in the baseline of the YFP/CFP ration measurements was detrended by calculating linear fit to the data using linear regression in GraphPad Prism. The data were normalized by ratio calculated from the respective fit and used for plotting in all figures.

