## Supplementary Figures for "Response of *Escherichia coli* chemotaxis pathway to pyrimidine deoxyribonucleosides"

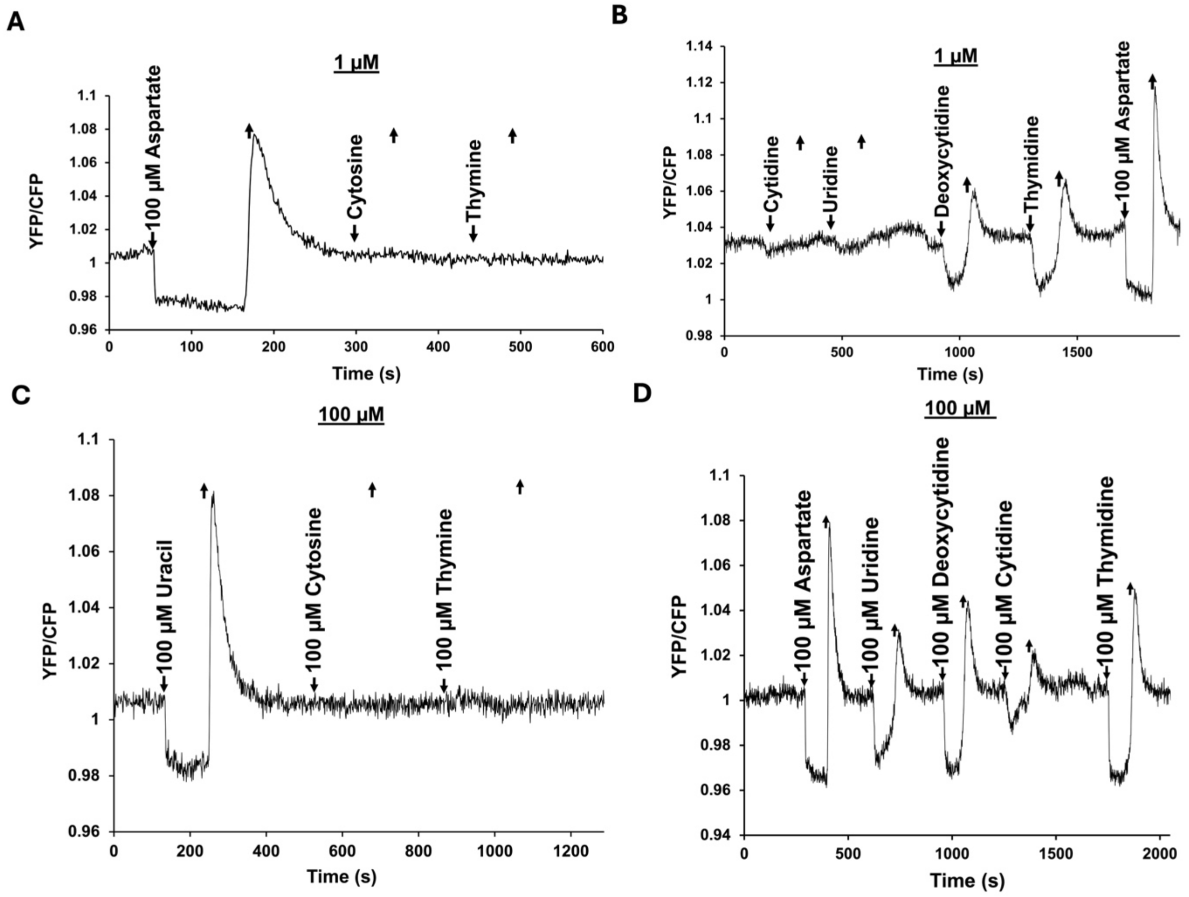


**FIG S1** FRET measurement of the chemotaxis pathway response to different concentrations of pyrimidine nucleosides and nucleobases. (A-B) Response to indicated compounds at 1 μM concentration. (C-D) Response to indicated compounds at 100 μM concentration. Measurements were performed as in Figure 1. Saturating stimulation with 100 μM L-aspartate was used as a control.

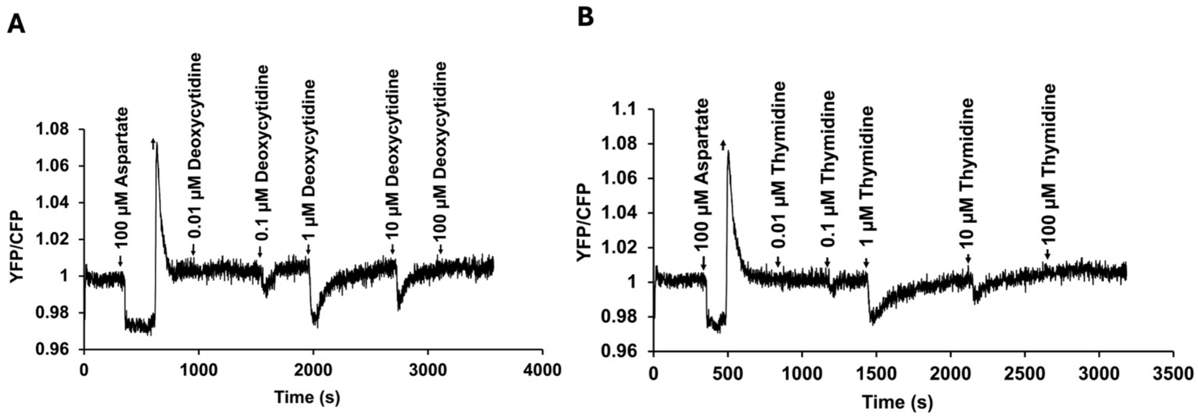


**FIG S2** Measurement of the dynamic range of the chemotactic response to deoxyribonucleosides. Concentration of deoxycytidine (A) or thymidine (B) was raised in 10-fold steps, and cells were allowed to adapt prior to each subsequent stimulation.

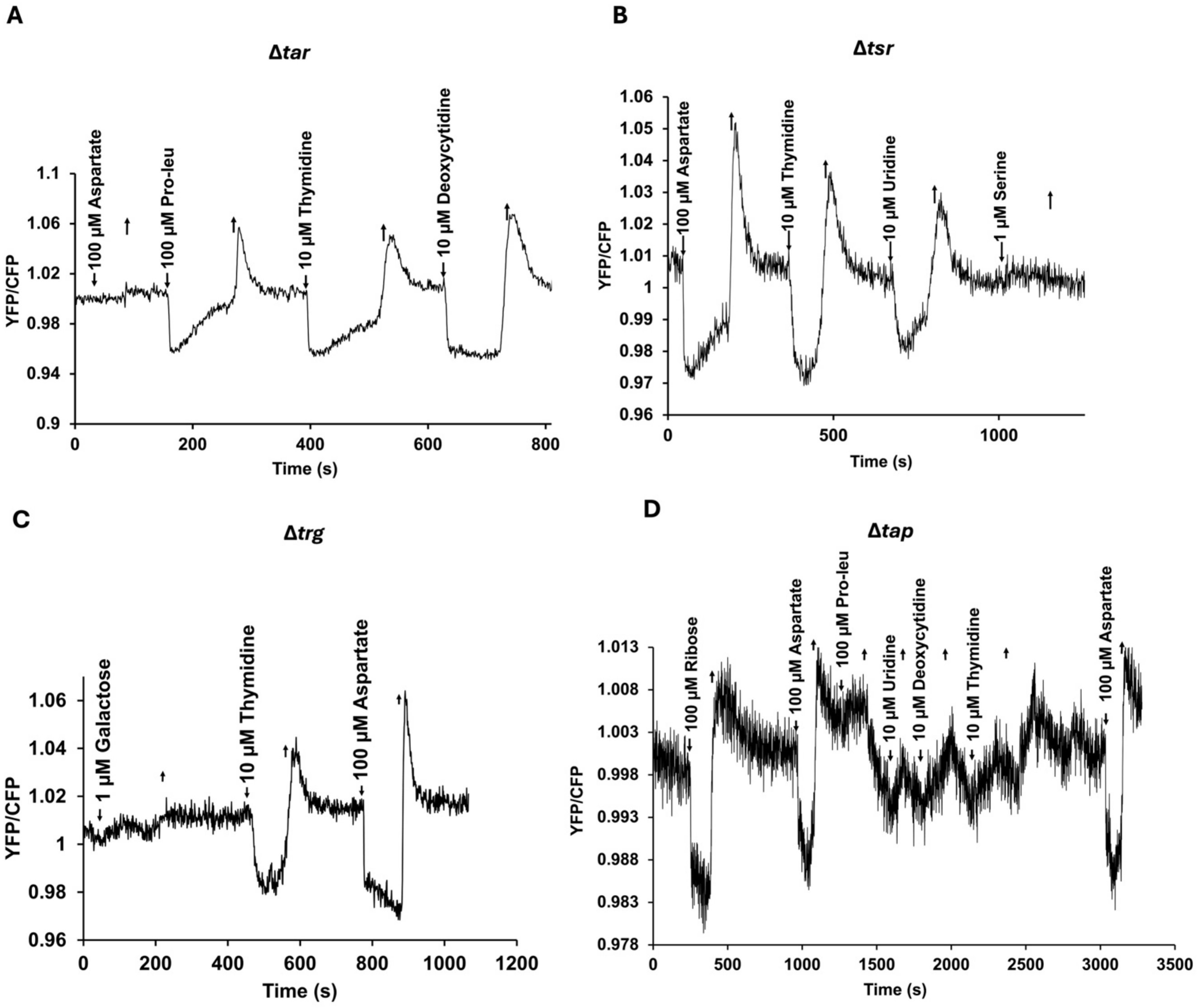


**FIG S3** Responses of Δ*tar* (A), Δ*tsr* (B), Δ*trg* (C), and Δ*tap* (D) chemoreceptor deletion strains to deoxyribonucleosides and control compounds at indicated concentrations. Measurements were performed as in Figure 1.

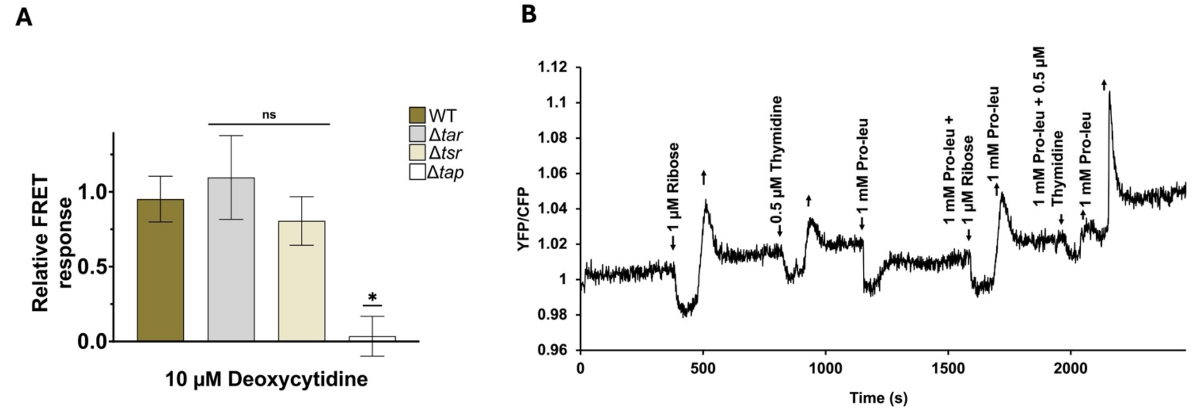


**FIG S4** Dependence of the response to deoxycytidine on Tap (A) and competition between proline-leucine (Pro-leu) and thymidine (B). Measurements were performed and quantified as described in Figure 2. Statistical significance was calculated using Student's two-sample *t*-test between wild- type and knockout (ns: non-significant, * *P*-value ≤0.05).
